# Group A *Streptococcus* Induces Lysosomal Dysfunction in THP-1 Macrophages

**DOI:** 10.1101/2022.06.17.496523

**Authors:** Scott T. Nishioka, Joshua Snipper, Jimin Lee, Joshua Schapiro, Robert Z. Zhang, Hyewon Abe, Andreas Till, Cheryl Y.M. Okumura

## Abstract

The human-specific bacterial pathogen Group A *Streptococcus* (GAS) is a significant cause of morbidity and mortality. Macrophages are important to control GAS infection, but previous data indicate that GAS can persist in macrophages. In this study, we detail the molecular mechanisms by which GAS survives in THP-1 macrophages. Our fluorescence microscopy studies demonstrate that GAS are readily phagocytosed by macrophages, but persist within phagolysosomes. These phagolysosomes are not acidified, which is in agreement with our findings that GAS cannot survive in low pH environments. We find that the secreted pore-forming toxin Streptolysin O (SLO) perforates the phagolysosomal membrane, allowing leakage of not only protons, but large proteins including the lysosomal protease cathepsin B. Additionally, GAS blocks the activity of vacuolar ATPase (v-ATPase) to prevent acidification of the phagolysosome. Thus, while GAS does not inhibit fusion of the lysosome with the phagosome, it has multiple mechanisms to prevent proper phagolysosome function, allowing for persistence of the bacteria within the macrophage. This has important implications for not only the initial response but the overall functionality of the macrophages, which may lead to the resulting pathologies in GAS infection. Our data suggests that therapies aimed at improving macrophage function may positively impact patient outcomes in GAS infection.

## 1 Introduction

GAS is a significant cause of morbidity and mortality worldwide, but especially in medically underserved areas where antibiotic therapy may not be promptly available. Infection produces wide-ranging clinical manifestations including pharyngitis, acute rheumatic fever (ARF) often leading to rheumatic heart disease, and invasive disease such as necrotizing fasciitis (1). Acting as the first line of defense against such infections, macrophages are critical for the early control and resolution of GAS infection (2,3). However, GAS has been shown to survive intracellularly in macrophages during both acute invasive soft tissue infection and asymptomatic carriage (3–5). Although GAS remains highly sensitive to beta-lactam antibiotics, the gold standard of treatment (6), the ability of bacteria to persist in macrophages and re-emerge after antibiotic treatment poses a therapeutic challenge (4).

As both phagocytes and antigen-presenting cells, macrophages function at the intersection of innate and adaptive immunity and are therefore critical for identifying and defending against infection. Macrophages engulf bacterial pathogens into phagosomes, which fuse with lysosomes that deliver proteolytic enzymes to facilitate bacterial destruction. These enzymes have optimal proteolytic activity under acidic conditions, and therefore lysosomal acidification is important for macrophage bactericidal function and subsequent antigen presentation (8). Inhibition of either lysosomal fusion with bacteria-containing phagosomes or phagolysosomal acidification is an evasion tactic employed by several bacterial pathogens including *Mycobacterium tuberculosis, Legionella pneumophila*, and *Staphylococcus aureus* (8–10). GAS may employ similar tactics to survive intracellularly within macrophages and to evade the adaptive immune response upon host re-infection, which may contribute to the development of pathologies such as toxic shock syndrome (11).

GAS is a human-specific pathogen and has evolved mechanisms to survive the immune response (12). Previous data have indicated that phagocytosed GAS can prevent fusion with destructive organelles such as azurophilic granules and lysosomes (13,14). Other reports have shown that while GAS-containing phagosomes do fuse with lysosomes, GAS survives within macrophages via the secretion of virulence factors such as Streptolysin O (SLO) and NADase (15,16). SLO is a cholesterol-dependent cytolysin that oligomerizes to form large pores (~30 nm) within the host cell membrane (17) that allows the loss of proteins up to 15nm in diameter (18). As a result, SLO is hypothesized to permeabilize the phagolysosomal membrane, permitting free flow of protons out of the phagolysosome and limiting the activity of lysosomal proteases that facilitate intracellular killing (16,19). Interestingly, it has also been reported that GAS is capable of preventing lysosomal acidification by preventing the recruitment of the vacuolar ATPase (v-ATPase) (20). In the context of these data, it is unclear what mechanisms are contributing to GAS intracellular survival in macrophages. In this paper, we used the human monocytic cell line THP-1 differentiated into macrophages to understand the molecular mechanism by which GAS survives in macrophages. We find that not only does GAS induce leakage of large proteins from the phagolysosome into the cytosol, but that acidification of the phagolysosome is also limited. This has important consequences for the survival of the bacteria within the macrophage and underscores the need to mount a proper macrophage response in GAS infection to improve patient outcomes.

## 2 Materials and Methods

### 2.1 Antibodies and chemicals

Antibodies to the following proteins were used in this study: EEA-1 (Abcam, ab2900), LAMP-2 (Abcam, ab25631), V0D1 (Abcam, ab56441), Cathepsin B (Cell Signaling Technology, D1C7Y), LAMP-1 (Cell Signaling Technology, #9091), V1A (Abnova, H00000523-A01), and beta-actin (Thermo Fisher Scientific, MA5-15739). Fluorescent secondary antibodies were purchased from Thermo Fisher Scientific. The following fluorescent probes were purchased from Thermo Fisher Scientific: Alexa Fluor 488 Hydrazide (A10436), Oregon Green 488 Anionic, Lysine Fixable Dextran,10,000 MW (D7171) and 70,000 MW (D7173), and LysoTracker DeepRed (L12492). 40,000 and 70,000 MW Fluorescein isothiocyanate (FITC) dextrans were purchased from Millipore Sigma. To generate antibody-coupled beads, 0.1um Carboxylate-Modified Microspheres (Thermo Fisher Scientific) were coupled to approximately 400mg normal pooled human serum (Thermo Fisher Scientific) using 2% 1-Ethyl-3-(3-dimethylaminopropyl) carbodiimide hydrochloride (QBiosciences). The reaction was quenched with 40mM ethanolamine and resuspended in PBS. Successful coupling of the beads was assessed by fluorescence microscopy showing increased phagocytosis by THP-1 macrophages compared with uncoupled beads (data not shown). 1mM L-leucyl L-leucine O-methyl ester (LLOMe, Cayman Chemical, #16008) was incubated with cells to induce lysosomal damage.

### 2.2 Bacterial strains

Wild-type (WT) GAS strain M1T1 5448 (M1 GAS) was originally isolated from a patient with necrotizing fasciitis and toxic shock syndrome (21). Isogenic mutant strains lacking Streptolysin O (ΔSLO), Streptolysin S (via lack of SagA, ΔSagA) and Emm1 (ΔM1) were previously described (22– 24). Bacterial strains were cultivated in Todd-Hewitt broth (THB) at 37°C. For all experiments, bacteria were grown to log phase in the presence of 1:200 pooled normal human serum (Thermo Fisher Scientific) to opsonize bacteria. The gene for monomeric WASABI (mWASABI) (7,25) was codon-optimized for expression in bacteria and synthesized in a shuttle vector (GenScript; see supplemental data for amino acid sequence). The gene was subcloned into pDCerm (26) and the resulting plasmid (pWASABI) was transformed into WT M1 GAS and the ΔSLO mutant. Plasmid-bearing strains were maintained in THB supplemented with 5ug/mL erythromycin. Heat-killed (HK) bacteria were prepared by incubating a known concentration of bacteria at 95°C for 10 min. followed by a 15 min. opsonization in 1:200 pooled normal human serum at room temperature.

### 2.3 Cell Culture

THP-1 cells were purchased from Sigma (cat. 88081201) and cultured in RPMI supplemented with 10% heat inactivated fetal bovine serum (FBS) (Corning), 2mM L-glutamine and 100 U/mL penicillin/100 µg/mL streptomycin at 37°C/5% CO2. Cells were differentiated to macrophages using 20nM phorbol 12-myristate 13-acetate (PMA) (MilliporeSigma) 24-48 hours prior to experiments.

### 2.4 Immunofluorescence

2.5× 10^5^ THP-1 cells were seeded on coverslips in the presence of 20nM PMA 24-48 hours prior to experiments. For dextran leakage assays, cells were incubated in media containing 20ug/mL of the indicated dextran or 500ug/mL 70kD dextran overnight, washed and chased in cell culture media for 2 hours prior to bacterial infection. For infection experiments with two rounds of infection, secondary infection bacteria were labeled with 10uM CellTracker Orange (Thermo Fisher Scientific) 30 min. prior to infection. Bacteria were combined with cells at an MOI = 10 in RPMI supplemented with 2% FBS only (no antibiotics) and plates were centrifuged at 500 x g to synchronize bacterial contact with cells. At the indicated time points, cells were fixed with 4% paraformaldehyde or 3.7% formaldehyde for 10 min. at room temperature. Cells were incubated with blocking solution (10% goat serum, 3% BSA and 0.1% Triton X-100 in PBS), then incubated with the indicated primary antibodies for 1 hour at room temperature in block solution. Cells were washed and incubated with the indicated secondary antibodies for 1 hour at room temperature. GAS opsonized in human serum were detected using fluorescent anti-human IgG antibodies where indicated. Cells were washed and mounted onto slides with ProLong antifade reagent with DAPI (Thermo Fisher Scientific). Slides were imaged on an inverted Leica TCS SP5 confocal microscope using a 63x/1.40 oil objective with 2-3x digital zoom at calibrated magnifications and recorded with LAS AF software (Leica). Quantitation of dextran and LAMP-1 colocalization with bacteria was analyzed with CellProfiler 3.1.9 (27) using the Otsu thresholding method and the MeasureImageOverlap module. To automatically count bacteria and cells, either fluorescence (bacteria) or brightfield (THP-1 macrophages) images were thresholded, objects were filled and average object size was used to count cells. For LAMP-1, colocalization is expressed as the percentage of GAS fluorescence that colocalized with LAMP-1 fluorescence, and the number of bacteria per cell was estimated by dividing the total number of bacteria by the total number of cells counted in each image. For dextran staining, colocalization is expressed as the percentage of dextran fluorescence that colocalized with GAS fluorescence. For quantitation of mWASABI-expressing bacteria within phagosomes, images were coded and counted manually in a blind fashion by at least 3 independent reviewers. For manually counted cells, the number of bacteria per cell and the number of infected cells was calculated by averaging the number of bacteria in infected cells across all counted images within an experiment. For all imaging data, at least 3 independent experiments were performed, at least 16 images containing >10 cells per image were analyzed and >100 cells were counted per time point.

### 2.5 Verification of pH sensitivity of mWASABI in GAS

To assess the fluorescence intensity of mWASABI in live GAS, log phase cultures of bacteria were incubated on coverslips in THB with 10mM HEPES adjusted to the indicated pH for 1 hour at 37°C. Bacteria were fixed with 2% paraformaldehyde and analyzed by confocal microscopy. Gain settings were adjusted to bacteria incubated in pH 7 media, and the settings were applied to all other conditions. Images were thresholded and the corrected total cell fluorescence (integrated density – average background integrated density) of each bacterial cell was measured using ImageJ.

### 2.6 Acidification assay

10^5^ THP-1 cells were seeded with 20nM PMA in 96-well black plates with clear bottoms overnight. The following day, cells were fed 500ug/mL 70kD FITC-dextran overnight. On the third day, cells were chased with normal cell culture media (RPM1 supplemented with 10% FBS, no antibiotics) for 2 hours prior to infection. Cells were infected at an MOI of 10 with either the indicated bacterial strain or Ab-conjugated latex beads in RPMI supplemented with 2% FBS only (no antibiotics) and plates were centrifuged at 500 x g to synchronize bacterial contact with cells. Fluorescence intensity at 480nm/535nm (Em/Ex) was monitored at 37°C in a fluorescent plate reader for 3 hours.

### 2.7 Bacterial survival in different pH media

Bacteria were incubated in THB with 10mM HEPES adjusted to the indicated pH. For bacterial growth, cultures were adjusted to an OD600nm of 0.1 and incubated at 37°C. OD600nm was measured every 15 min. For bacterial survival, 2 × 10^5^ cfu log phase bacteria were added to 250ul pH-adjusted culture media in 96 well plates. At the indicated time point, a 25ul aliquot was removed from the culture and quantitated by plating on agar plates.

### 2.8 Cell fractionation and Western blot

Following a 30 min. infection period, a total of 10^7^ PMA-differentiated cells were scraped and collected in 1 mL of fractionation buffer (50 mM KCl, 90 mM K-Gluconate, 1 mM EGTA, 5 mM MgCl2, 50 mM sucrose, 20 mM HEPES, pH 7.4, 5 mM glucose, 1X HALT phosphatase/protease inhibitor cocktail (Thermo Fisher Scientific), 1ug/mL pepstatin (Millipore Sigma), and 1mM PMSF (Millipore Sigma)). Cells were lysed by nitrogen cavitation equilibrated on ice for 30 min. at 400 psi. The resulting lysates were fractionated into membrane and cytosolic fractions by centrifugation at 16,000 x g at 4°C for 15 min. (pellet = membrane fraction, supernatant = cytosolic fraction). Protein concentrations were measured by BCA assay (Thermo Fisher Scientific). 10ug of each sample was run on SDS-PAGE gels and transferred to a 0.2um PVDF membrane. Blots were blocked with 5% (w/v) non-fat dry milk in 1X Tris-buffered saline with 0.1% Tween 20 (TBST). Blots were probed with the indicated antibodies overnight at 4°C. Relative protein concentrations were quantified by densitometry analysis in ImageLab v. 6.0.1 (BioRad). v-ATPase assembly was measured by normalizing V1A densitometry values in the membrane fraction to the V0D1 loading controls in membrane fractions and calculating the ratio of V1A in the sample compared with uninfected macrophages (28).

### 2.9 Cathepsin B activity assay

4 × 10^6^ PMA-differentiated THP-1 cells were infected at an MOI = 10 for 60 min. followed by lysis with 30ug/mL digitonin in acetate buffer (50 mM Na-acetate pH 5.6, 150 mM NaCl, 0.5 mM EDTA) and protease inhibitors (1X HALT phosphatase/protease inhibitor cocktail (Thermo Fisher Scientific), 100 μM PMSF, 1 μg/mL pepstatin)) (29). The cell lysate was separated into cytosolic and membrane fractions as described above. Cathepsin B activity in cytosolic fractions was measured using the SensoLyte 520 Cathepsin B Assay Kit (AnaSpec) according to the manufacturer’s instructions. Relative fluorescence units were normalized to the corresponding protein concentration for each sample and compared with the uninfected cells. Data from 3 independent experiments were combined.

### 2.10 v-ATPase activity assay

Cell membrane fractions containing lysosomes were prepared using 30ug/mL digitonin in acetate buffer as described above. 10ug of each sample was equilibrated in 200ul assay buffer (10mM HEPES, 5mM MgCl2, 125mM KCl, pH = 7.0) (30) at room temperature for 30 min. in the presence or absence of 200nM bafilomycin (MilliporeSigma). 1mM fresh ATP in 200mM Tris pH = 7 was added to each sample and incubated at room temperature for 2.5 hours. 25ul of the reaction was collected and diluted in ultrapure water, and liberation of free phosphate was measured using the Malachite Green Phosphate Assay Kit according to the manufacturer’s instructions (MilliporeSigma). Corrected absorbance values (= A_sample_ - A_substrate blank_ - A_buffer blank_) were compared with a phosphate standard curve to determine phosphate concentration. v-ATPase activity was calculated as the difference in phosphate concentration between samples incubated with and without bafilomycin.

### 2.11 Statistical analysis

Growth curve data was analyzed using the Growthcurver package in R (31) and comparison of the intrinsic growth rate (r) and carrying capacity (K) was analyzed by one-way ANOVA and Tukey’s post-hoc test. All other data were analyzed using Prism v. 9.3.1 (GraphPad Software), on arcsin transformed data where indicated, by one-way ANOVA with Tukey’s (normally distributed data) or Dunnett’s or Kruskal-Wallis (nonparametric data) multiple comparison tests. Data in Figures 2D, 2G and Supp. Figure 4B were also analyzed by 2-way ANOVA to ensure day to day experimentation and/or individual counters did not contribute to the differences observed in the data (data not shown). Outliers were assessed by the ROUT method (Q = 1%). For all data presented, ****P<0.0001, ***P<0.001, *P<0.05, n.s., not significant.

## 3 Results

### 3.1 Bacterial trafficking in THP-1 macrophages

Because there are reports that GAS can prevent (14) or allow (16) lysosomal fusion upon phagocytosis, we first examined the intracellular trafficking of GAS in THP-1 macrophages. Bacterial contact with cells was initiated by centrifugation to synchronize macrophage uptake of bacteria and phagolysosomal maturation was monitored by immunofluorescence in a short time course infection experiment (Fig. 1, 2). In agreement with the literature (14,16,32), bacteria were rapidly engulfed by THP-1 cells by our earliest measured time point at 7.5 minutes (Fig. 1B, 2D) and colocalized with the early phagosomal marker EEA-1 (Fig. 2C, D). Interestingly, bacteria in phagosomes also began colocalizing with the lysosomal marker LAMP-1 as early as 7.5 minutes (Fig. 1B). Maximal colocalization with LAMP-1 began at 30 minutes and GAS remained within phagolysosomes throughout the later time points (Fig. 1A, B). Surprisingly, but in agreement with other reports (32), the number of bacteria per cell was relatively constant throughout the experiment (Fig. 1C). These data indicated that bacteria were trafficked as expected to phagolysosomes, but bacteria were held in these compartments without observable bacterial elimination or replication within this time frame. Although we synchronized the infection, there was a possibility that this result was due to continuous infection of macrophages and turnover of bacteria. To address this, we performed the trafficking experiment with a second round of bacterial infection (Fig. 1D). Bacteria introduced in the secondary infection were rarely found in macrophages and were mostly extracellular (Fig. 1D), in agreement with a report for *Salmonella* Typhimurium that the frequency of macrophage reinfection after the primary infection is rare (33). Thus, bacteria from the primary infection persist in THP-1 macrophage phagolysosomes.

**Figure 1:**
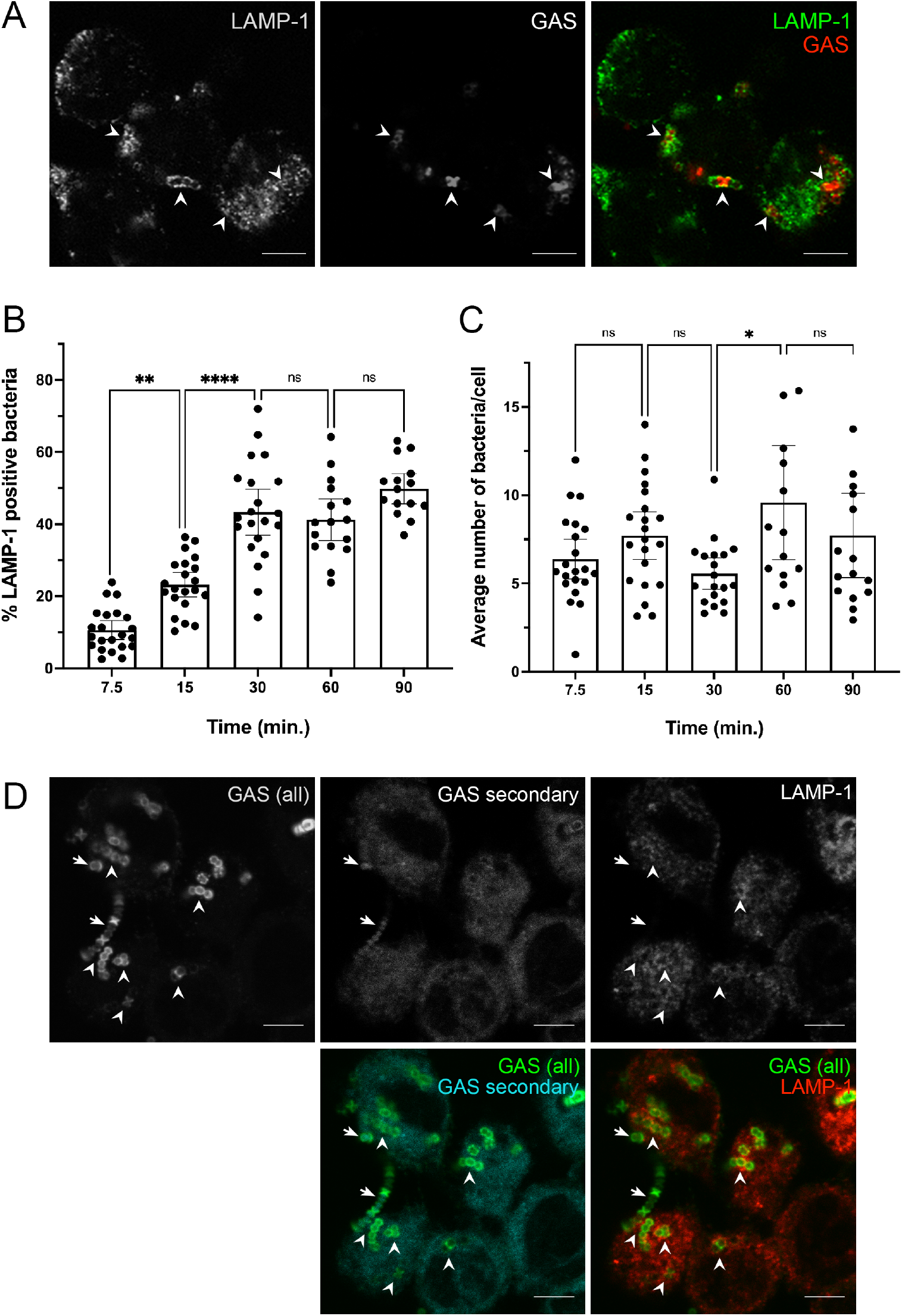
GAS persists in phagolysosomes of THP-1 cells. **(A**,**B)** THP-1 cells were infected with GAS for the indicated times, fixed and probed with anti-LAMP-1 and secondary fluorescent antibodies (green). Bacteria opsonized with human serum were detected with anti-human IgG antibody (red). **(A)** Representative image of LAMP-1 colocalized with GAS in THP-1 cells at 60 min. post-infection. Arrowheads indicate examples of colocalization. Scale bar = 5µm. **(B)** Quantitation of LAMP-1 positive bacteria at the indicated time points. **(C)** Average number of bacteria per cell at the indicated time points. **(D)** Representative image of THP-1 cells exposed to a primary (green only) GAS infection for 30 min. followed by a secondary (cyan and green) infection for 30 min. (60 min. total infection time). LAMP-1 staining is indicated in red. Arrowheads indicate examples of colocalization between primary infecting GAS and LAMP-1. Arrows indicate GAS from the secondary infection. Scale bar = 5µm. For all graphs, >100 cells were counted, data from three independent experiments were combined and results are given as mean ± 95% CI. Data were analyzed on arcsin transformed data by one-way ANOVA with Tukey’s multiple comparisons test.

**Figure 2:**
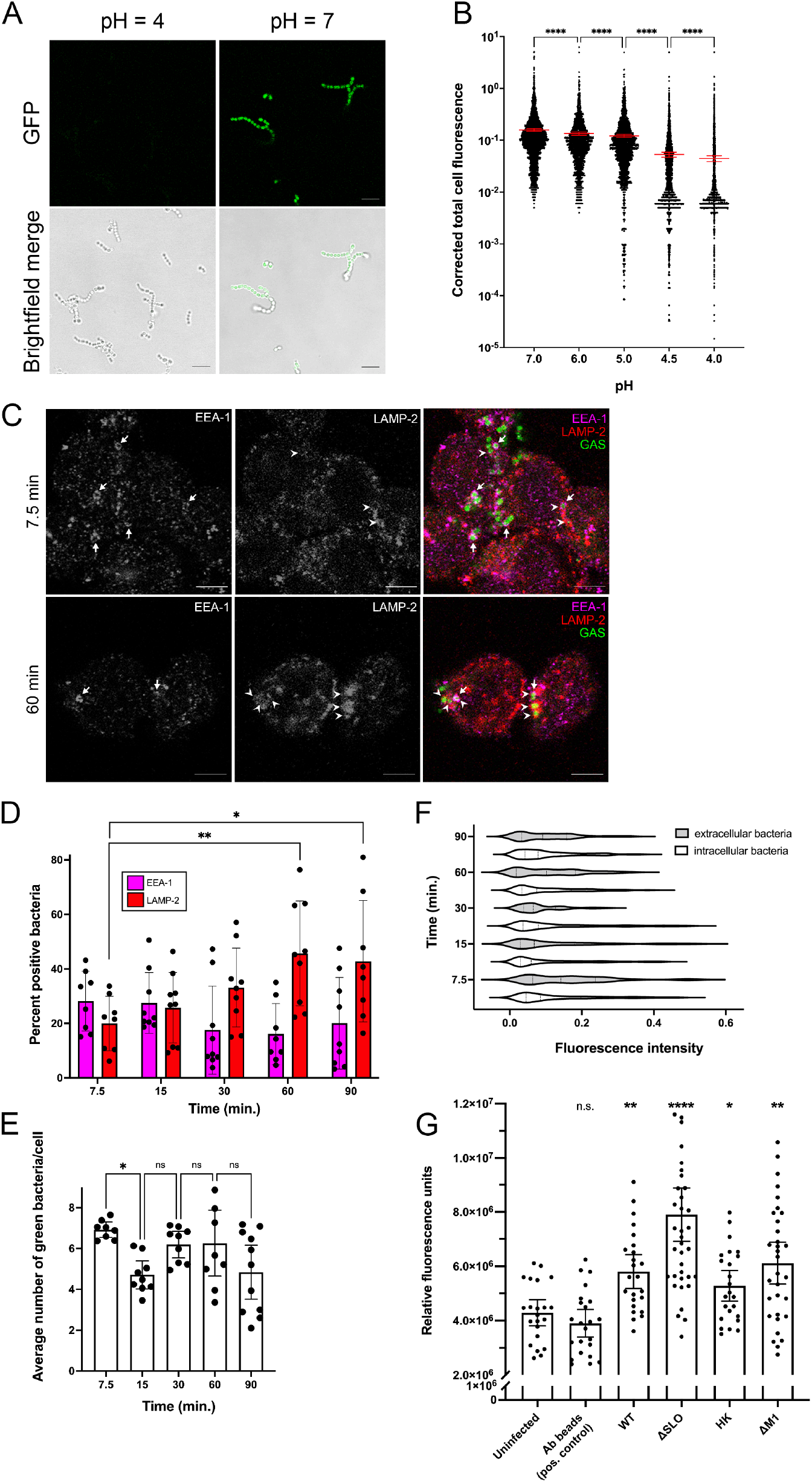
GAS persists in non-acidified phagolysosomes of THP-1 cells. **(A)** GAS expressing mWASABI were incubated in media at the indicated pH for 1 hour. Representative green fluorescence (mWASABI) and brightfield images are shown. Scale bar = 5µm. **(B)** Corrected total cell fluorescence of green fluorescence signal from bacteria expressing mWASABI incubated in media at the indicated pH for 1 hour. **(C)** Representative fluorescence microscopy images of GAS-infected THP-1 cells at 7.5 and 60 min. post-infection. Arrows denote examples of bacteria encapsulated in early phagosomes (EEA-1, magenta), arrowheads denote examples of bacteria encapsulated in phagolysosomes (LAMP-2, red). All images were taken with a 63x objective with 2x digital zoom, scale bar = 5µm. **(D)** Quantitation of bacteria colocalized with phagosomes (EEA-1) or phagolysosomes (LAMP-2) at the indicated time points. **(E)** Average number of bacteria per cell at the indicated time points. **(F)** Violin plot of fluorescence signal intensity from intracellular (clear bars) or extracellular (shaded bars) bacteria expressing mWASABI. **(G)** Fluorescence signal from cells fed 70kD FITC dextran and infected with immunoglobulin-conjugated beads (Ab beads), the indicated GAS strain or heat-killed (HK) bacteria was measured by plate reader. Statistical differences were calculated by comparing samples to uninfected cells. Data from at least three independent experiments were combined. For (D) and (E), >100 cells were counted, data from at least 3 individual counters of at least three independent experiments are shown. Results are given as mean ± 95% CI and statistics were performed on arcsin transformed data by one-way ANOVA with Dunnett’s multiple comparisons test.

### 3.2 GAS reside in non-acidified phagolysosomes in THP-1 macrophages

Acidification is critical for optimal activity of the proteolytic enzymes in the lysosome (34). Because bacteria appeared to be held in phagolysosomes without observable elimination, we wondered whether phagolysosomes containing GAS were acidified. Lysotracker is an acidotropic dye commonly used to track acidic compartments such as lysosomes in the cell. However, because GAS is an acid-producing bacterial species, we tested whether Lysotracker would be an accurate indicator of lysosomal pH in GAS-infected cells. Although Lysotracker colocalized with lysosomal markers in THP-1 cells, the staining and uptake was inconsistent (Supp. Fig. 1A). Furthermore, bacteria were brightly stained with Lysotracker in both infected cells and in the absence of cells (Supp. Fig. 1A-C). This made it difficult to determine whether and to what extent phagolysosomes containing GAS were acidified. We therefore expressed a pH-sensitive green fluorescent protein variant mWASABI (7) in GAS to monitor phagolysosomal pH of GAS-infected cells (Fig. 2A). Green fluorescence produced by live bacteria was appropriately quenched in low pH environments (Fig. 2A, B). Notably, pH was distinguishable by fluorescence intensity between pH 5.0 and 4.5 (Fig. 2B).

We infected THP-1 macrophages with mWASABI-expressing GAS and performed a detailed trafficking assay of GAS through the phagolysosomal pathway (Fig. 2C, D). For this experiment, we used the lysosomal protein LAMP-2 to confirm the lysosomal fusion we observed earlier (Fig. 1) since there is conflicting evidence in the literature that GAS prevents lysosomal fusion (13). Because bacteria were tracked by mWASABI expression instead of immunofluorescent staining colocalization with phagosomal markers was difficult (Fig. 2C, D). To quantitate the data, we manually counted bacteria that colocalized with phagolysosomal markers in a blinded manner (Fig. 2D). We again found that GAS was initially located in phagosomes, as indicated by EEA-1 staining, which was followed by rapid bacterial colocalization with the mature lysosomal marker LAMP-2 within the first 15 minutes of infection (Fig. 2C, D), similar to the colocalization dynamics observed with LAMP-1 staining (Fig. 1). An average of 50% of the counted intracellular bacteria were colocalized with LAMP-2 at the 60 and 90 minute post-infection time points, which was significantly increased compared with the 7.5 minute time point (Fig. 2C, D). Although there were some fluctuations in the number of bacteria in phagolysosomes (Fig. 2E), the number of intracellular bacteria remained relatively constant at all time points, consistent with our initial data (Fig. 1C).

Supporting this data, the percentage of infected cells was similar across all time points (Supp. Fig. 2). This corroborated our finding that little to no bacterial eradication or replication was occurring within this time frame (Fig. 2E). Surprisingly, we also found that the green fluorescence signal did not diminish over time, indicating that the intracellular bacteria were not in an acidic environment (Fig. 2C, F). Fluorescence signal intensity of intracellular bacteria in phagolysosomes was not significantly different than that of the extracellular bacteria in our images (Fig. 2F). As the lysosomal lumen is typically between pH 4.5–5 when acidified (35), a pH level that our mWASABI probe can differentiate (Fig. 2A, B), these data indicated that GAS were maintained in phagolysosomes that are not appropriately acidified.

We confirmed that GAS-infected phagosomes were not acidified with a second method using pH-sensitive FITC-conjugated 70kD dextran particles. Acidification of the lysosomes resulted in quenching of the fluorescence signal in uninfected and control cells infected with immunoglobulin-conjugated beads (Fig. 2G). However, the fluorescence signal was increased in cells infected with all tested strains of GAS, indicating no pH change (Fig. 2G). The low magnitude of difference we observed in our GAS-infected cells compared with the uninfected cells may be due to the limited ability of cells to uptake such a large molecule (70kD) which produced variation between experiments (Fig. 2G). Nevertheless, these results confirmed that GAS-infected phagolysosomes are not properly acidified. Additionally, there are differences in signal between GAS strains (Fig. 2G), but all cells infected with GAS show increased fluorescence signal compared with the uninfected cells. This ability of GAS to prevent lysosomal acidification may account for the persistence of the bacteria in THP-1 cells (Fig. 1C, 2E).

### 3.3 GAS does not tolerate acidic lysosomal conditions

GAS is a lactic acid-producing bacterial species and possesses systems such as the arginine deiminase (ADI) pathway that confer acid tolerance (36,37). As expected, we found that the final pH of both buffered and unbuffered culture medium after bacterial growth was acidic (Table 1). However, in a physically constrained environment such as the lysosome, the effectiveness of such systems may be limited. The pH of the lysosomal lumen has been reported to range between pH 4.5-5 (35). We wondered whether a pH similar to that found in the lysosome adversely affected bacterial growth and survival. In timed growth experiments, bacteria grown in buffered media adjusted to pH 7 had no significant difference in maximum population size, but a statistically significant increase in the intrinsic growth rate of the population compared with unbuffered media (Fig. 3A). Bacteria grown in media adjusted to pH 6 grew at a significantly slower rate and did not reach a similar maximum population size in the time frame of our experiment compared with unbuffered media (Fig. 3A). Bacteria grown in media adjusted to pH 5 or lower did not demonstrate measurable growth (Fig. 3A). Thus, GAS growth is sensitive to decreasing pH and is inhibited at a pH found in the lysosomal environment.

**Table 1:** Final pH of media after bacterial growth, mean ± SD.

| starting pH | 4.0 | 5.0 | 6.0 | 7.0 | unbuffered THB<br>(6.6) |
| --- | --- | --- | --- | --- | --- |
| ending pH | 4.10 ± 0.02 | 5.07 ± 0.02 | 4.99 ± 0.01 | 5.23 ± 0.08 | 5.63 ± 0.06 |

**Figure 3:**
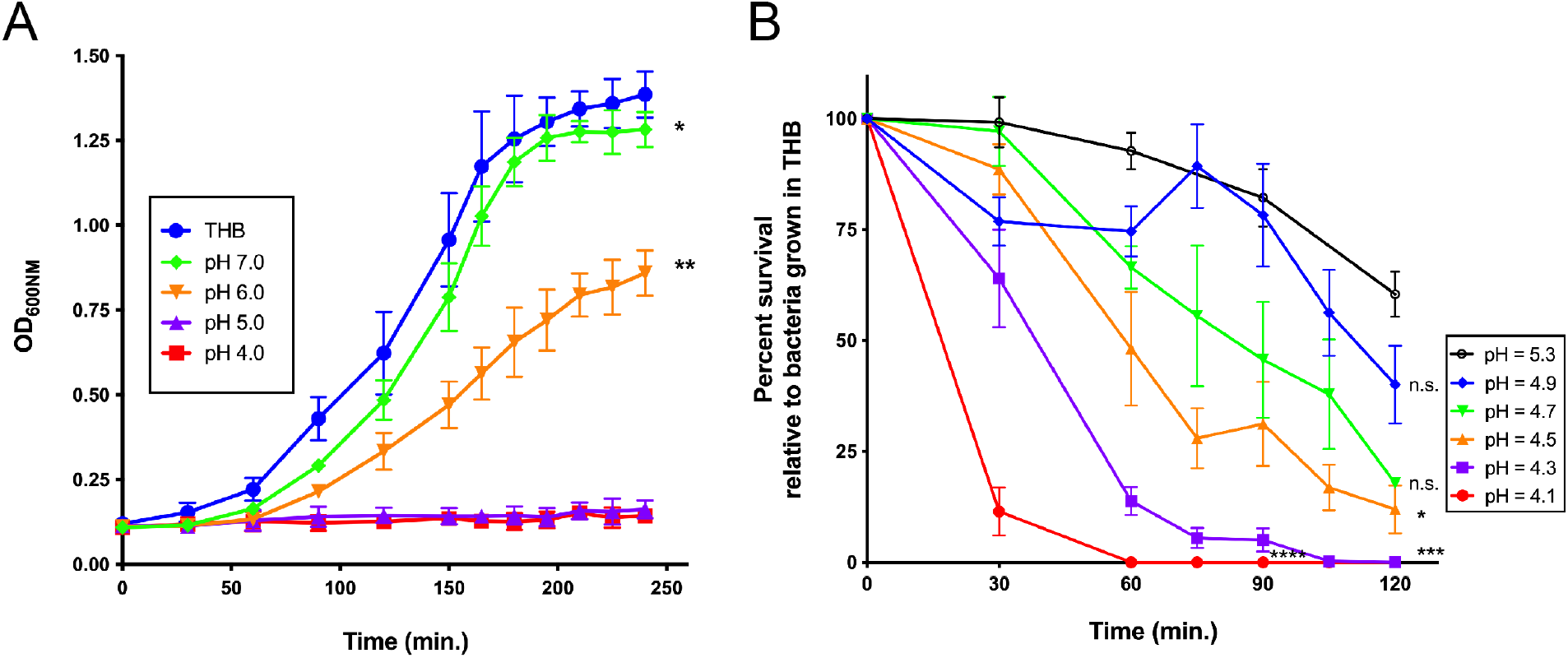
GAS does not survive in low pH environments. **(A)** Growth curves of bacteria grown in buffered media or unbuffered bacterial media (THB) at the indicated pH. Results are given as mean ± 95% CI. Statistics in (A) indicate differences in intrinsic growth rate compared with bacteria grown in unbuffered media tested by one-way ANOVA with Tukey’s multiple comparisons test. **(B)** Bacterial survival in buffered media at the indicated pH compared with survival of bacteria in unbuffered bacterial media (THB). Statistics in (B) indicate differences in area under the curve compared with bacteria grown in pH = 5.3 media tested by one-way ANOVA with Tukey’s multiple comparisons test. For each experiment, samples were prepared in triplicate and data from at least three independent experiments were combined and results are given as mean ± SEM.

The final pH of the stationary phase cultures in both buffered and unbuffered culture media (Table 1) indicated that although GAS may not replicate in acidic conditions, it may be capable of survival. Quantitative plating of cultures at various time points throughout the growth experiment indicated that bacteria in pH 5 media were present (data not shown), but not measurably growing (Fig. 3A). We therefore performed a detailed time course to monitor bacterial survival in low pH media (Fig. 3B). Bacteria incubated in unbuffered media did not significantly grow during the time frame of the survival experiment (data not shown). Similar to other studies (38,39), we found that GAS does not survive in low pH media (Fig. 3B). Although bacteria incubated in up to pH 5.3 media could survive for a short time, bacterial numbers declined at later time points (Fig. 3B), in agreement with pH 5 being the limit at which bacteria can survive (37,40) and the final pH of stationary phase bacterial cultures (Table 1). Our data affirm that GAS cannot survive in the acidified lysosome and therefore, limiting or preventing acidification (Fig. 2) is a survival strategy that GAS may use to persist in phagolysosomes.

### 3.4 Streptolysin O creates large perforations in the phagolysosome

Although GAS has several acid stress response strategies (41), a better tactic may be to avoid the acidification of the phagolysosome altogether. The inability of bacteria to survive in low pH environments (Fig. 3) led us to examine whether GAS infection has a mechanism to prevent phagolysosomal acidification. Pore-forming toxins such as SLO create pores as large as 25-30nm in diameter (17). Others have shown that pore-forming toxins, including SLO, allow escape of hydrogen ions that prevent acidification of the lysosome (16,42). Streptolysin S (SLS) can similarly perforate membranes (43). We fed THP-1 macrophages fluorescent-conjugated molecules and dextrans of various sizes to assess pore size and leakage from phagolysosomes in GAS-infected cells. The ability to measure phagosomal leakage was confirmed using the lysosomal permeabilization agent LLOMe (Supp. Fig. 3). LLOMe permeabilizes lysosomes to molecules up to at least 4.4kD, but not to molecules greater than 10kD (29). Retention of the 10kD, but not the 570Da probe in LLOMe-treated cells confirmed uptake of the probe in phagolysosomes and appropriate release into the cytosol upon permeabilization (Supp. Fig. 3A). Cells loaded with probes of varying sizes were infected with GAS (Fig. 4A, B). As expected, WT GAS caused probes up to 40kD in size, approximately 9nm in diameter (44), to leak from phagolysosomes (Fig. 4A, B). We next used heat-killed bacteria (HK) that retain surface structure but are unable to secrete proteins such as SLO and SLS. We found that the fluorescent probes were retained and colocalized with heat-killed bacteria (Fig. 4A, B), suggesting that a secreted toxin is responsible for leakage. To determine which toxins were responsible for this leakage, we infected cells with bacteria lacking different proteins. ΔSLO GAS-infected cells retained the probes, while cells infected with ΔSagA GAS (lacking SLS) allowed leakage of the probes, similar to the WT strain (Fig. 4A, B). In our assays, SLS did not significantly damage the phagolysosomal membrane, as indicated by the lack of leakage of even the small 570Da probe (Supp. Fig. 3B). Mutant bacteria lacking the surface protein M1 (ΔM1), but with unaltered expression of SLO and SLS behaved similarly to WT strains (Fig. 4A, B). Our data further indicate that SLO allowed probes up to 70kD in size, approximately 13nm in diameter (44), to escape from the lysosomes (Fig. 4B), in agreement with the reported pore size for SLO (17). These data indicate that SLO is the primary pore-forming toxin that perforates the phagolysosome and allows leakage of not only protons, but relatively large proteins.

Because such large molecules can leak from GAS-infected phagosomes, we next tested whether lysosomal enzymes could escape into the cytosol through SLO-mediated pores. We infected cells with WT or ΔSLO GAS and collected membrane and cytosolic cell fractions. Lack of lysosomal markers in the cytosolic fraction indicated little to no contamination of the cytosolic fraction with lysosomes (Fig. 4C). The membrane fraction contained most of the lysosomal enzyme cathepsin B as expected (Fig. 4C). However, we noted a small, but visible increase in the amount of cathepsin B present in the cytosolic fraction of WT-infected cells compared with uninfected and ΔSLO-infected cells (Fig. 4C). In order to verify that cathepsin B was present in the cytosol, we measured cathepsin B activity in the cytosolic fraction. Cytosolic fractions from WT-infected cells contained the highest amount of cathepsin B activity compared with uninfected cells (Fig. 4D). Cytosolic fractions from ΔSLO-infected cells (Fig. 4B) did not exhibit cathepsin B activity higher than uninfected cells (Fig. 4D). These data confirmed that SLO secreted by WT bacteria perforates the phagolysosomal membrane and allows the leakage of large proteins such as the lysosomal enzyme cathepsin B into the cytosol. This loss of proteolytic enzymes from the phagolysosome in addition to the loss of protons likely contributes to the ability of GAS to persist in the phagolysosome during infection.

**Figure 4:**
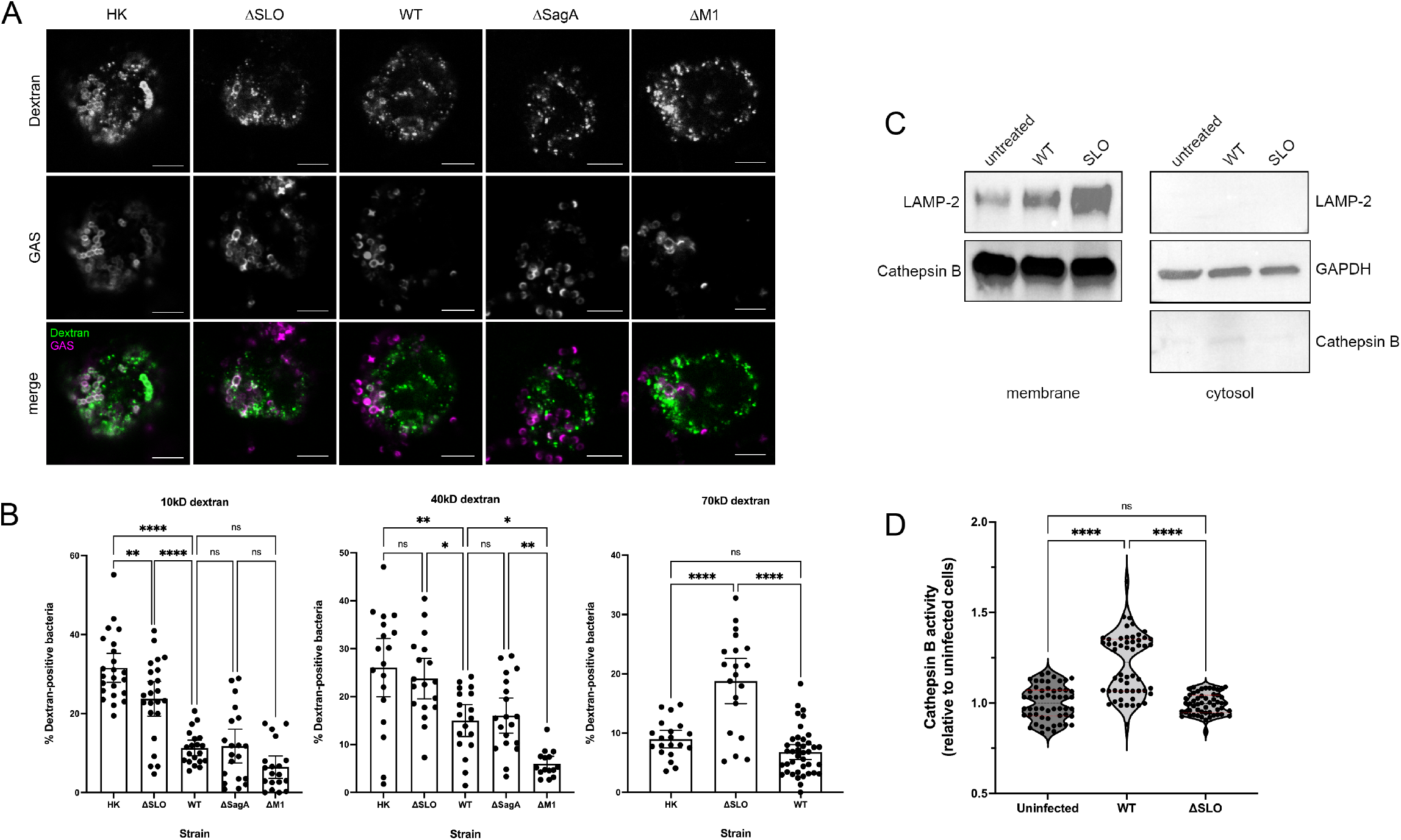
SLO induces phagolysosomal perforation and protein leakage. **(A)** Representative images of the 40kD dextran probe (top panels, green in merged images) with indicated bacterial strains (middle panels, magenta in merged images). All images were taken with a 63x objective with 2x digital zoom, scale bar = 5µm. **(B)** Quantitation of colocalization of indicated bacterial strains with 10, 40 and 70kD dextran probes, >100 cells were counted. **(C)** Cells were infected with the indicated bacterial strains, fractionated into membrane (left) and cytosolic (right) fractions, and probed with the indicated antibodies: LAMP-2 (lysosomal marker; loading control for membrane fraction), GAPDH (cytosolic marker; loading control for cytosolic fraction) and cathepsin B (lysosomal enzyme). **(D)** Cathepsin B activity was measured in cytosolic cell fractions collected from cells infected with the indicated strains. Data were normalized to the corresponding protein concentration for each sample and compared with the uninfected cells. Data in (B) and (D) are combined from at least three independent experiments. Results are given as mean ± 95% CI and and statistics were performed on arcsin transformed data by one-way ANOVA with Tukey’s multiple comparison test (B) or one-way ANOVA with Kruskal-Wallis multiple comparison test (D).

### 3.5 GAS prevents proper function of vacuolar ATPase

Although SLO-mediated pores could cause leakage of protons from the phagolysosome, reports in the literature also suggest that the v-ATPase that is required for acidification of the lysosome is non-functional in GAS-infected cells (20). Supporting this, we found that the phagolysosomes of macrophages infected with ΔSLO and heat-killed bacteria were also not acidified (Fig. 2G). Furthermore, ΔSLO bacteria are trafficked to phagolysosomes, maintain the fluorescent signal of mWASABI, and are maintained at constant numbers in THP-1 cells similar to WT bacteria (Supp. Fig. 4). These data are consistent with the idea that phagolysosomes infected with ΔSLO bacteria are also not acidified. We therefore investigated whether v-ATPase is properly assembled and functional in GAS-infected cells. v-ATPase consists of the membrane-bound F0 subunit (containing the V0D1 protein) that must assemble with the cytosolic F1 subunit (containing the V1A protein) for proper function (28). We monitored v-ATPase assembly by tracking movement of the V1A subunit from the cytosolic to membrane fractions. Cells fed Ig-conjugated beads (Ab beads) were included as a positive control (Fig. 5A, B). We observed no difference in v-ATPase subunit assembly in cells infected with GAS compared to Ab bead-fed or uninfected cells (Fig. 5A, B), indicating that the acidification machinery was recruited properly to phagolysosomes. However, when we measured v-ATPase activity in membrane fractions of GAS-infected cells, we found significantly decreased v-ATPase activity in fractions from cells infected with all strains of GAS compared with uninfected cells (Fig. 5C). These data are in agreement with the lack of acidification in GAS-infected cells observed in our experiments using pH-sensitive dextrans (Fig. 2G). Therefore, though the v-ATPase is properly assembled in GAS-infected cells, the activity of this enzyme is limited, preventing proper acidification of the phagolysosome. Thus, our data suggests that GAS persists in macrophages by employing a multi-pronged approach to avoid the acidification and destructive mechanisms of the phagolysosome.

**Figure 5:**
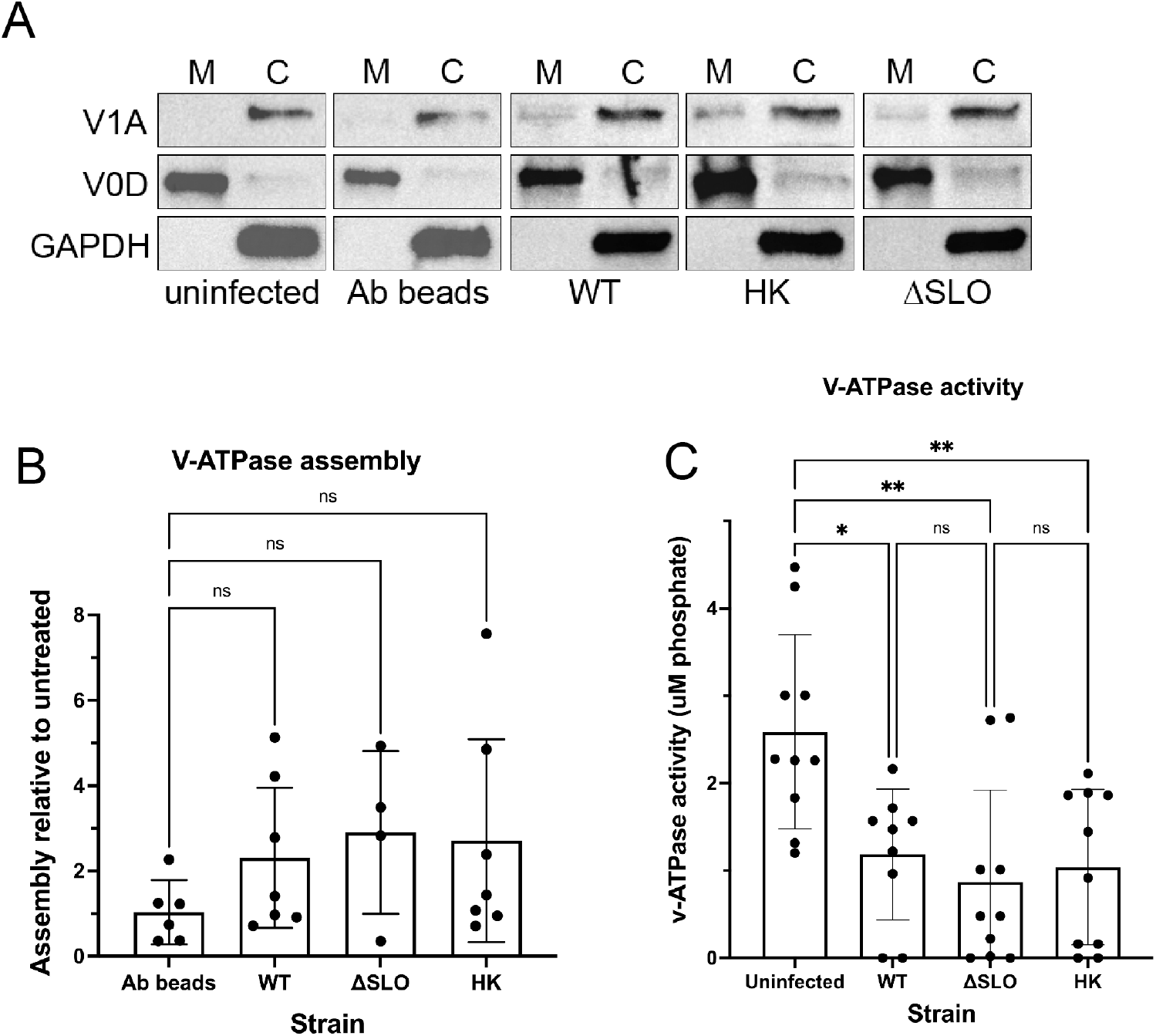
GAS infection limits v-ATPase activity. **(A)** Cells were uninfected, infected with immunoglobulin-conjugated beads (Ab beads, control), the indicated GAS strain or heat-killed (HK) bacteria and fractionated into membrane (M) and cytosolic (C) fractions. Fractions were probed with antibodies to V1A, V0D or GAPDH proteins. Representative data are shown. **(B)** Quantitation of data shown in (A). V1A band intensities were quantified and normalized to V0D (membrane) or GAPDH (cytosol). Relative assembly was calculated as the amount of V1A in the membrane fraction of the indicated infected sample compared with the uninfected sample for each experiment. **(C)** v-ATPase activity assay in membrane fractions from cells infected with the indicated GAS strain. v-ATPase activity was calculated by determining the concentration of free phosphate (released by the ATPase) of the samples and subtracting the concentration of free phosphate in samples incubated with bafilomycin (v-ATPase inhibitor) from the samples incubated without bafilomycin. Data in (B) and (C) are combined from at least three independent experiments and results are given as mean ± 95% CI and analyzed by one-way ANOVA with Tukey’s multiple comparison test.

## 4 Discussion

The human-specific pathogen GAS has co-evolved with the human immune system, and has multiple mechanisms for survival within the host (1,12). Although GAS is an extracellular bacterial pathogen, phagocytic cells such as macrophages can readily engulf bacteria (14,16,32). The ability of GAS to not only survive in macrophages, but to potentially alter their function, escape downstream immune responses, and be sheltered from antibiotic treatment provides a basis for successful persistence in humans.

In our experiments, we found that GAS was readily phagocytosed and phagosomes quickly fused with lysosomes, but that GAS remained in these compartments (Fig. 1, 2). There is conflicting evidence in the literature about whether lysosomes fuse with GAS-containing phagosomes in macrophages, which may be due to a difference in cell origin or source of antibodies (14,16). We find that in THP-1 macrophages, lysosomal fusion with GAS-containing phagosomes occurs, as confirmed by colocalization with two independent lysosomal antibodies (Fig. 1, 2). We recognize that although our studies may be limited due to the exclusive use of cell culture, our data are in agreement with several studies regarding lysosomal fusion with GAS-infected phagosomes (16,32). In addition, we corroborated previous data that SLO induces perforation and damage of the phagolysosomal membrane (Fig. 4) (16,45) which promotes bacterial survival. In addition to loss of lysosomal enzymes and protons, we also demonstrated that GAS limits phagolysosomal acidification to increase bacterial survival (Fig. 2, 5). Although ΔSLO GAS also prevented phagolysosomal acidification (Fig. 2, 5), others have shown that ΔSLO GAS are less fit for survival in macrophages and other cells (16,46). ΔSLO GAS is likely still susceptible to other macrophage killing mechanisms such as reactive oxygen species or xenophagy (46). Finally, our data demonstrate that bacteria persist in phagolysosomes without replication or escape into the cytosol (Fig. 1, 2, Supp. Fig. 4) (14,16).

This is in agreement with another study of GAS in THP-1 cells (32), but different from what has previously been observed in epithelial and U-937 cells, where GAS escapes into the cytosol and replicates (15,32,45). In our experiments, not all GAS were in LAMP-positive compartments (Fig. 1, 2), which may be due to lysophagic clearance of GAS-damaged lysosomal membranes (32,47). However, the downstream fate of GAS in non-functional lysosomes in THP-1 macrophages remains to be elucidated.

The pH of the lysosomal lumen has been reported to range between pH 4.5-5 (35). However, we found that acidified medium was sufficient to kill GAS (Fig. 3). These results are somewhat surprising since not only does GAS produce lactic acid, but previous reports suggest GAS has multiple mechanisms for acid tolerance, including a F0/F1 proton pump and the arginine deiminase pathway which produces alkali (36,37,41,48). However, there may be a limit to which these systems work, especially within an enclosed environment such as in vitro conditions or in the phagolysosome, as demonstrated by the threshold for bacterial survival in low pH medium (Fig. 3) (37). Additionally, GAS lacks enzymes such as glutamate decarboxylase and urease (41), making neutralization of highly acidified environments a less viable strategy. Therefore, two mechanisms to prevent acidification, perforating the phagolysosomal membrane (Fig. 4) and blocking v-ATPase activity (Fig. 5) provide a means to survive within the phagolysosomal compartment. Other bacteria such as *Legionella pneumophila* have proteins such as SidK, which binds v-ATPase subunits to prevent activity (30). Our data indicate that neither SLO nor any other secreted factor is responsible for interfering with v-ATPase activity (Fig. 5C), and the identity of the GAS protein(s) responsible for inhibiting v-ATPase activity remains to be elucidated.

Previous data and our results indicate SLO causes leakage of lysosomal contents (Fig. 4). These proteins include active lysosomal enzymes such as cathepsin B (Fig. 4). The cathepsin B activity we measured was subtle, likely because the enzyme optimally functions in a low pH environment (8), which is neither achieved in GAS-infected lysosomes (Fig. 2, 5) nor in the cytosol. However, releasing active lysosomal proteases could have downstream effects on the cell such as activation of the NLRP3 inflammasome and pyroptosis (49). Furthermore, SLO promotes macrophage apoptosis by releasing SLO from the lysosome and causing mitochondrial damage (22,50). Other proteins released from the infected lysosome could also include bacterial proteases such as SpeB, whose presence in the cytosol could lead to NLRP3 inflammasome activation and release of the proinflammatory cytokine IL-1β (51–55). Persistent IL-1β activation as a result of GAS infection has been linked with ARF and rheumatic heart disease (56). Thus, besides enabling bacterial survival and blunting the normal macrophage response, this alternative macrophage response may lead to pathologies such as ARF that are observed during GAS infection. Studies have shown that macrophages are crucial for GAS clearance (2), but the loss of macrophage function may also have detrimental effects on the long-term response to GAS infection, including the inability to generate protective antibodies. Therapies aimed at improving macrophage function may therefore improve GAS infection outcomes. Although SLO is a promising target, our work shows that additional drugs to restore v-ATPase activity would be necessary to fully restore macrophage function. Restoration of macrophage function in GAS infection would allow individuals to mount a proper adaptive immune response to confer protective immunity.

## Supporting information

Supplemental data and SFig. 1-4

## 6 Author Contributions

SN, JS, AT and CO contributed to conception and design of the study. All authors performed the experiments and analyzed data. CO wrote the first draft of the manuscript. All authors contributed to manuscript revision, read, and approved the submitted version.

## 8 Acknowledgments

We thank Cheldon Alcantara and Ryan Hino for contributions to experiments, Karen Molinder for helpful discussions and Kelvin Zheng, Jessica Kim, Stephanie Peacock, Isabel Perez, Kevin Kim, and the Fall 2021 Bio 395 GRR course for contributions to data analysis in this paper.

