## Supplemental data and SFig. 1-4 for "Group A *Streptococcus* Induces Lysosomal Dysfunction in THP-1 Macrophages"

### *Supplementary Material*

#### 1 Supplementary Data

mWASABI amino acid sequence:

MVSKGEETTMGVIKPDMKIKLKMEGNVNGHAFVIEGEGEGKPYDGTNTINLEVKEGAPLPF
SYDILTTAFSYGNRAFTKYPDDIPNYFKQSFPEGYSWERTMTFEDKGIVKVKSDISMEEDSFI
YEIHLKGENFPPNGPVMQKETTGWDASTERMYVRDGVLKGDVKMKLLLEGGGHHRVDFK
TIYRAKKAVKLPDYHFVDHRIEILNHDKDYNKVTVYEIAVARNSTDGMDELYK

#### 2 Supplementary Figures

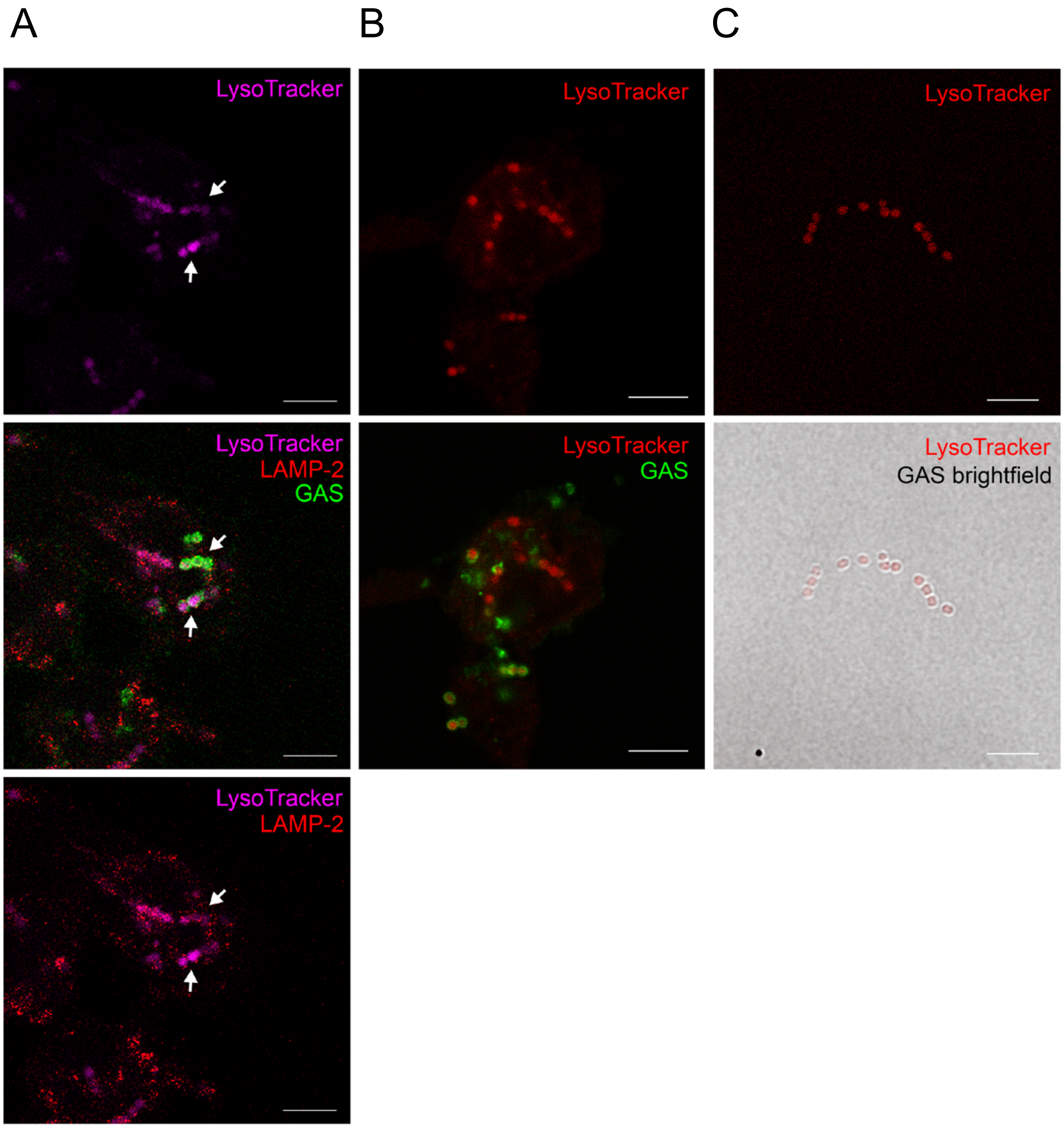


**Supplemental Figure 1: Lysotracker detects GAS.** Cells were loaded with 100nM LysoTracker Deep Red (Thermo Fisher Scientific), then infected with GAS for 30 min. Cells were fixed and stained with antibodies to LAMP-2 (lysosome) or anti-human IgG antibodies to detect opsonized GAS as indicated: **(A)** LysoTracker (magenta), LAMP-2 (red), and GAS (green). Arrows indicate examples of colocalization between GAS and LysoTracker. **(B)** Lysotracker (red) and GAS (green). **(C)** Bacteria alone incubated with 100nM Lysotracker (red). All images were taken with a 63x objective with 2x digital zoom, scale bar = 5μM.

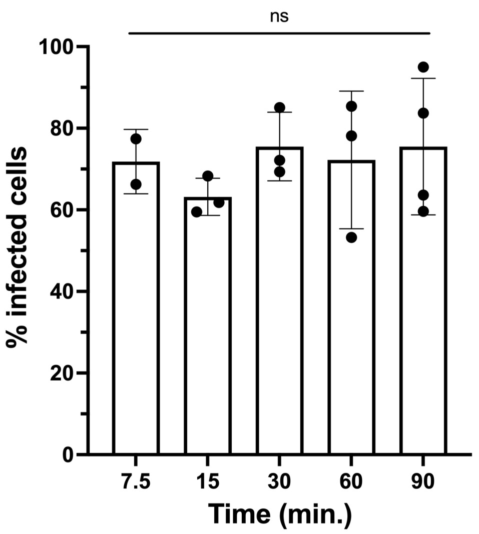


**Supplemental Figure 2: Percentage of cells infected with GAS.** For each time point, >100 cells were counted, data from at least 2 individual counters of independent experiments are shown. Results are given as mean ± 95% CI and statistics were performed on arcsin transformed data by one-way ANOVA with Tukey's multiple comparisons test.

.

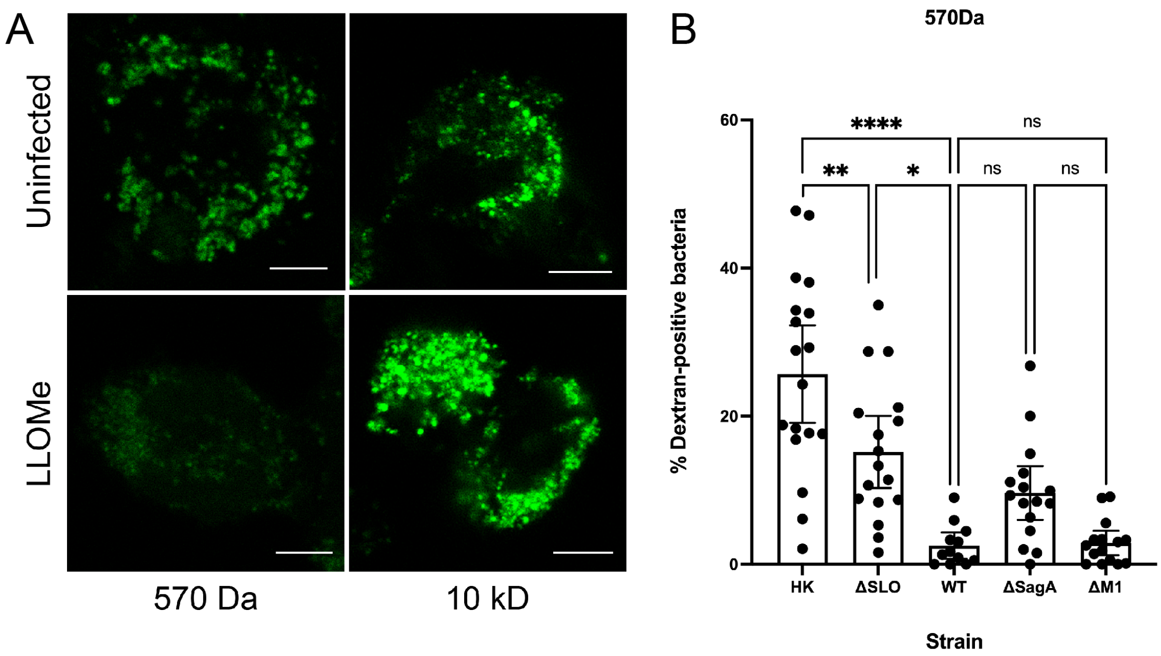


**Supplemental Figure 3: Fluorescent probes appropriately monitor phagolysosomal leakage.** Cells were loaded with Alexa-fluor 488 dye (570 Da) or 10kD Oregon Green dextran (10kD) for 24 hrs., then either uninfected or treated with 1mM LLOMe. **(A)** Representative images of fluorescent probe-loaded cells in the presence or absence of LLOMe. All images were taken with a 63x objective with 2x digital zoom, scale bar = 5μM. **(B)** Quantitation of colocalization of Alexa-fluor 488 (570Da) with indicated bacterial strains. Data from three independent experiments were combined and results are shown as mean ± 95% CI. Statistics were performed on arcsin transformed data by one-way ANOVA with Tukey's multiple comparisons test.

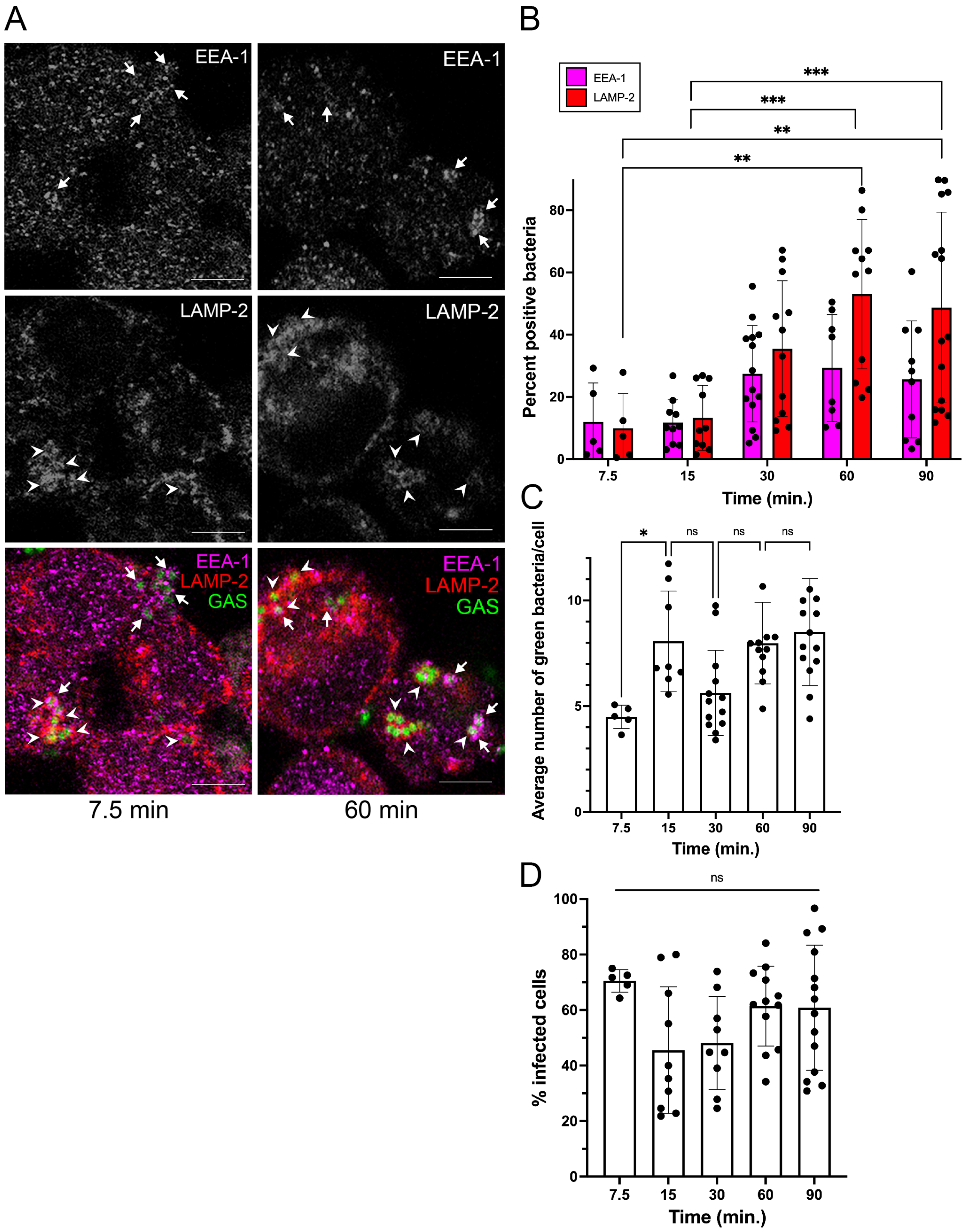


**Supplemental Figure 4: ΔSLO GAS persist in THP-1 phagolysosomes. (A)** Representative fluorescence microscopy images of ΔSLO-infected THP-1 cells at 7.5 (left column) and 60 min. (right column) post-infection. Arrows denote examples of ΔSLO bacteria (green) encapsulated in early phagosomes (EEA-1, top panel and magenta in image overlay), arrowheads denote examples of ΔSLO bacteria (green) encapsulated in phagolysosomes (LAMP-2, middle panel and red in image overlay). All images were taken with a 63x objective with 2x digital zoom, scale bar = 5µM. **(B)** Quantitation of bacteria colocalized with phagosomes (EEA-1) or phagolysosomes (LAMP-2) at the indicated time points. **(C)** Average number of bacteria per cell at the indicated time points. **(D)** Percentage of cells infected with GAS. For (B-D) >100 cells were counted, data from at least 3 individual counters of at least three independent experiments are shown. Results are given as mean ± 95% CI and statistics were performed on arcsin transformed data by one-way ANOVA with Dunnett's (B) or Tukey's (C, D) multiple comparisons test.
